# Preservation of memory B cell homeostasis in an individual producing broadly neutralising antibodies against HIV-1

**DOI:** 10.1101/2024.02.05.578789

**Authors:** Sarah Griffith, Luke Muir, Ondrej Suchanek, Joshua Hope, Corinna Pade, Joseph M Gibbons, Zewen Kelvin Tuong, Audrey Fung, Emma Touizer, Chloe Rees-Spear, Andrea Nans, Chloe Roustan, Yilmaz Alguel, Douglas Fink, Chloe Orkin, Jane Deayton, Jane Anderson, Ravindra K Gupta, Katie J Doores, Peter Cherepanov, Áine McKnight, Menna Clatworthy, Laura E McCoy

## Abstract

Immunological determinants favouring emergence of broadly neutralising antibodies are crucial to the development of HIV-1 vaccination strategies. Here, we combined RNAseq and B cell cloning approaches to isolate a broadly neutralising antibody (bnAb) ELC07 from an individual living with untreated HIV-1. Using single particle cryogenic electron microscopy (cryo-EM), we show that the antibody recognises a conformational epitope at the gp120-gp41 interface. ELC07 binds the closed state of the viral glycoprotein causing considerable perturbations to the gp41 trimer core structure. Phenotypic analysis of memory B cell subsets from the ELC07 bnAb donor revealed a lack of expected HIV-1-associated dysfunction, specifically no increase in CD21^-^/CD27^-^ cells was observed whilst the resting memory (CD21^+^/CD27^+^) population appeared preserved despite uncontrolled HIV-1 viraemia. Moreover, single cell transcriptomes of memory B cells from this bnAb donor showed a resting memory phenotype irrespective of the epitope they targeted or their ability to neutralise diverse strains of HIV-1. Strikingly, single memory B cells from the ELC07 bnAb donor were transcriptionally similar to memory B cells from HIV-negative individuals. Our results demonstrate that potent bnAbs can arise without the HIV-1-induced dysregulation of the memory B cell compartment and suggest that sufficient levels of antigenic stimulation with a strategically designed immunogen could be effective in HIV-negative vaccine recipients.

## Introduction

People living with HIV-1 can produce strain-specific neutralising antibodies (nAbs) that apply selection pressure on the virus, inevitably resulting in viral escape^1^. Given that the virus can easily escape these host nAbs, it is not surprising that they do not offer protection against re-infection with the many circulating strains of HIV-1^2^. However, after prolonged infection, a subset of individuals (10-30%) develop nAbs that exhibit cross-neutralisation of diverse viral strains, while a smaller proportion (1-10%) produce sera containing broadly neutralising antibodies (bnAbs) that are active across HIV-1 clades^3^. To date, HIV-1 vaccination attempts have induced largely strain-specific nAbs that have not proven effective in preventing infection^4,5^. By contrast, passive transfer of HIV-1 bnAbs provide protection in animal models^6,7^ alongside promising results using them as therapeutics^8^. However, it has not yet been possible to robustly induce bnAbs by vaccination. Moreover, it remains unclear why only a minority of individuals develop bnAbs during natural HIV-1 infection.

Many studies have explored the development of bnAbs, and although certain traits have been associated with HIV-1 neutralisation breadth, none appear to be solely responsible for or able to predict their emergence^9^. Multiple studies observed an association between the duration of HIV-1 infection and the acquisition of serum neutralisation breadth^10–13^, with bnAb lineages emerging as late as five years after HIV-1 exposure^14–17^. Given the correlation between neutralisation breadth and the duration of untreated HIV-1 infection, it is not surprising that an association with high viral load and quasi species diversity has been identified^10,12,18^. In addition to virological features, there are clear associations of neutralisation breadth with immune parameters. Thus, CD4+ T cell counts, which are the main cell type infected by HIV-1 and are depleted during untreated infection, are inversely associated with plasma neutralisation breadth, particularly at viral setpoint or early in infection^10,11,19,20^. However, the counts of circulating T_FH_ cells, CD4+ T cells crucial for antibody affinity maturation in the germinal centre (GC), were found to correlate with the development of neutralisation breadth in humans^21^ and primates^22^. Similarly, higher levels of the key GC-recruiting chemokine CXCL13 have been associated with neutralisation breadth^23–25^, suggesting that GC reactions are enhanced in individuals that develop bnAbs. Furthermore, a positive association has been found with high levels of expression of *RAB11FIP5* in natural killer (NK) cells and the development of neutralisation breadth^26^.

While conflicting results have been reported with donor-related parameters, including ethnicity and gender^10,12,27^, bnAbs have been identified in individuals with autoimmune diseases^28^. Concordantly, a number of bnAbs have been described as polyreactive or autoreactive^29^. The development of such bnAbs, therefore, implies a level of B cell dysregulation associated with escape from tolerance checks. To date, investigation into the phenotype of B cells associated with neutralisation breadth has frequently focused on characterisation of autoreactivity of individual antibodies and the differences in antibody repertoires between people living with HIV-1 based on their neutralisation capacity^30,31^. However, lower numbers of total B cells during early acute infection have been positively associated with neutralisation breadth, although crucially it was found that higher numbers of B cells specific for autologous HIV-1 envelope glycoprotein (Env) in this pool were also linked to bnAb development^32^. Although HIV-1 does not infect or replicate in B cells, the virus can have profound effects on the B cell compartment in people who are not on suppressive anti-retroviral therapy (ART)^33^. In particular, hyperactivation of B cells leads to a decreased proportion of naïve B cells and an increased proportion of both immature/transitional B cells and plasmablasts in circulation^33^. Combined with increased numbers of plasma cells, this gives rise to non-HIV-1-specific polyclonal hypergammaglobulinemia^34,35^. Moreover, HIV-1 infection can be associated with major alterations in the memory B cell subsets, classified by expression of CD27 and CD21 into resting memory (RM: CD27+ CD21+), activated memory (AM: CD27+ CD21-) and tissue-like memory (TLM; CD27-CD21-) B cells. RM are diminished and TLM B cells are expanded in untreated HIV-1 infection. Interestingly, TLM B cells display enhanced expression of inhibitory receptors that overlap with those of exhausted T cells, as well as homing receptors to inflammatory sites rather than the GC^36^. Investigation of HIV-1 Env-specific antibodies from TLM B cells revealed a lower level of somatic hypermutation (SHM) than antibodies from RM B cells^37^ suggesting restricted affinity maturation.

In this study, we combined single B cell cloning and transcriptomic analysis in an elite bnAb donor. Using cryo-EM, we characterised a potent bnAb clone derived from the patient, which revealed a novel conformational epitope at the gp41-gp120 interface. Characterisation of the memory B cells of this bnAb donor uncovered a striking preservation of RM B cells and no increase in TLM B cells. These findings are in sharp contrast to the profiles normally associated with HIV-1 viraemia across both total memory B cells and HIV-1 Env reactive memory B cells, suggesting that bnAbs can develop without the HIV-1-induced dysregulation of the memory B cell compartment that is commonly associated with viraemia.

## Results

### Identification of a bnAb ELC07 in an elite HIV-1 neutraliser from a historical cohort

To identify individuals with a broadly neutralising antibody response, we screened the East London cohort. All donors had been living with HIV-1 for more than a year and were not on ART, as they were recruited before 2010 and did not have an AIDS-defining illness or CD4+ cell count below 200 cells/mm^3^ in line with prior UK treatment guidelines^38^. This cohort included individuals from geographically diverse regions with different circulating HIV-1 clades^39^. The historical neutralisation data^39^ were examined, and individuals with no reactivity against the negative control virus and neutralisation of more than one tier 2 or 3 HIV-1 isolate with a 50% inhibitory dilution (ID_50_) titer > 100 were selected for in-depth characterisation against the 6-virus panel (Fig 1A)^10,40^. To assess the extent of neutralisation breadth, the ID_50_ titers against this previously validated indicator panel of 6 pseudotype viruses (PVs) were log-transformed and averaged to calculate a neutralisation score^10,40^. Plasma samples were ranked in order of their neutralisation scores to identify moderate neutralisers (score ≥ 0.5), broad neutralisers (score ≥ 1), and elite neutralisers (score ≥ 2)^10^ (Fig 1A). As previously described, the latter group had the highest potential to produce bnAbs ^41,42^. One elite neutraliser, T125, a donor with clade C HIV-1, exhibited remarkably potent neutralisation (ID_50_ titer >1000) of four PVs in the panel from three different clades, and consequently achieved the highest neutralisation score of 3.19 (Fig 1A). Plasma epitope mapping revealed more than a 3-fold change in neutralisation potency against N160A/K169T and N276D/N462D PV mutants indicating the presence of trimer apex and CD4 binding site (CD4bs) specific antibodies in the plasma (Fig S1).

**Figure 1:**
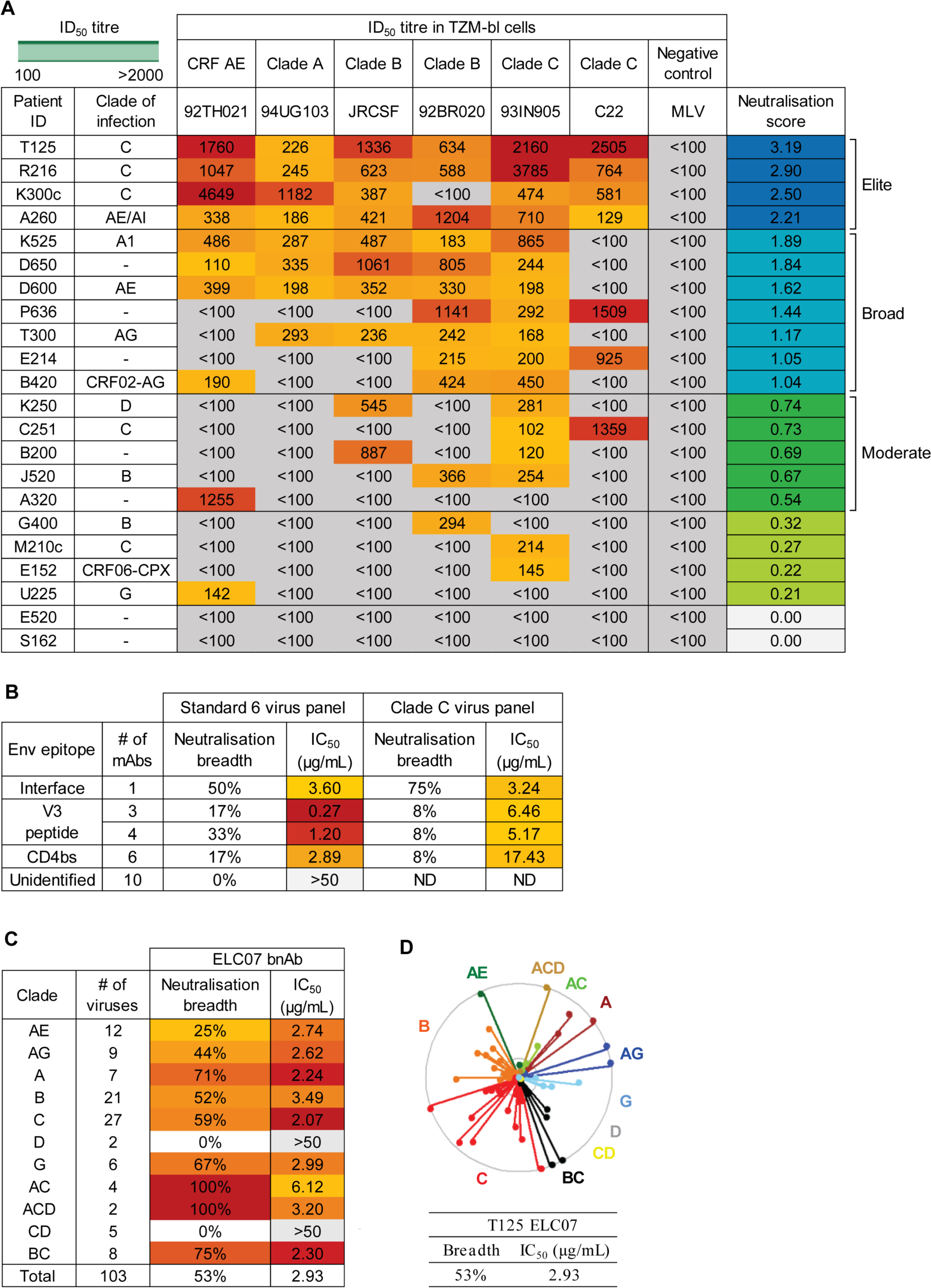
Identification of an elite neutraliser that produced an interface-targeting bnAb ELC07. (A) Plasma/serum neutralisation ID_50_ titers against a predicative standard 6 PV panel and a negative control (MLV) PV for patients in the East London cohort (n=22) with a colour gradient from yellow to red for lowest to highest respectively, organised by neutralisation score to identify elite neutralisers (score > 2), broad neutralisers (score >1) and moderate neutralisers (score > 0.5). (B) Percentage neutralisation breadth and median IC_50_ values for the viruses neutralised in the predicative standard 6 PV panel and standard 12 clade C PV panel, with mAbs grouped by the epitope targeted. (C) Percentage neutralisation breadth and geomean IC_50_ values by T125 bnAb ELC07 against viruses (n=103) from the standard multi-clade 118 PV panel. (D) Neutralisation dendrogram of IC_50_ values by T125 bnAb ELC07 against viruses (n=103) from the standard multi-clade 118 PV panel, coloured by the virus clade. The outer circle represents an IC_50_ of <1 µg/mL, the inner circle <5 µg/mL and the centre of the circle 50 µg/mL.

Single HIV-1 Env reactive memory B cells from T125, at two timepoints four months apart (at least one year after HIV-1 infection while ART naive), were then isolated by flow activated cell sorting (FACS) using streptavidin-conjugated stabilised HIV-1 Env trimers. The antibodies encoded by isolated B cells were cloned, produced recombinantly and tested for neutralisation. While the majority of the resulting patient-derived monoclonal antibodies (mAbs) had no neutralising activity, some V3 peptide- and CD4 binding site-specific mAbs were found to neutralise a limited number of PVs in the 6-virus panel and the standard clade C panel (Fig 1B). By contrast, one clone, designated ELC07, was able to neutralise 50% of the 6-virus panel and 75% of the clade C panel (Fig 1B). Further evaluation of ELC07 against 103 PVs demonstrated that this bnAb neutralised a wide variety of HIV-1 clades with a total breadth of 53% and relatively low median potency of 2.93 μg/mL (Fig 1C, D), similar to the breadth and potency of previously characterised gp120-gp41 interface targeting bnAbs^43–46^. Therefore, competition binding experiments were performed that revealed that ELC07 competed with a previously described interface bnAb 3BC315^45^ for binding to HIV-1 Env. Moreover, akin to 3BC315, ELC07 displayed a markedly enhanced neutralisation activity against HIV-1 Env carrying the T90A mutation, which abrogates glycosylation of Asn88 (Fig S1)^45^. By contrast, other characterised gp120-gp41 interface bnAbs, such as ASC202, depend on Asn88 glycosylation^46^.

### Structure of HIV-1 Env in complex with ELC07 bnAb

The competition with 3BC315 suggested that ELC07 belongs to the gp120-gp41 interface bnAbs, which display considerable heterogeny in their epitopes and effects on HIV-1 Env glycoprotein^45–48^. To determine the structural basis for the broad HIV-1 neutralisation activity of ELC07, we imaged the stabilised trimeric HIV-1 BG505 Env gp140 SOSIP.664 construct^49^ in the presence of near equimolar amounts of ELC07 Fab by cryo-EM. BG505 was one of the strains used to identify the ELC07 producer B cell and is well-suited for cryo-EM^50^. Classification of single particle images revealed the presence of HIV-1 Env trimers in complex with one or two Fab moieties bound, and the structure of the protein complex containing a single Fab was refined to a global resolution of 2.9 Å (Fig 2; Fig S2A). The local resolution of the reconstruction reached 2.5 Å throughout the core of the Env trimer and 2.5-2.8 Å within its interface with the antibody (Fig S2A). Both HIV-1 Env trimer, including 46 N-linked glycans, and the Fab molecule were well defined in the cryo-EM map (Fig S2B), allowing building and refinement of a high-quality atomistic model (Fig 2A; Table S1). ELC07 binds at the base of the HIV-1 Env trimer, primarily engaging one gp41 subunit, and making additional direct interactions with gp120 from the same gp120-gp41 protomer. The majority of the interactions involve the Cys-Cys loop of gp41 (spanning BG505 HIV-1 Env residues 582-628) and the heavy chain of the antibody. The ELC07 epitope centers on the invariant gp41 residue Trp623 (HXB2 Trp553), which forms an aromatic stacking interaction with Tyr98 from CDR H3. Tyr98 along with Arg100D and His100F (all located in CDR H3) engage in hydrophobic interactions with gp41 Ile603 (HXB2 Ile544) and Leu619 (HXB2 Trp549) as well as with gp120 Thr499 ((HXB2 Thr445). While Ile603 is invariant and Thr499 conserved in >95% HIV-1 strains, the position 619 is conserved as hydrophobic, although the BG505 strain is an exception. The unnatural residues Cys605 and Cys501 introduced to stabilise the Env trimer via a gp120-gp41 disulfide bond (SOS)^49^ are found at the periphery of the epitope where they contribute to the hydrophobic patch interacting with CDR H3 (Fig 2B). In the majority of HIV-1 strains, these positions are occupied by small hydrophobic residues (Ala and Thr, respectively), which are expected to engage in similar hydrophobic interactions with ELC07 CDR H3. Several residues from CDR H1 (Thr31), CDR H2 (Ile52, Leu53 and Val54) and CDR H3 (Ser99 and Phe100E) contribute to an extended hydrophobic groove at the tip of ELC07 Fab. While bound to BG505 Env, the groove accepts the side chain of Met535 (Met478 in HXB2) residing at the beginning of gp41 heptad repeat 1 (HR1), it appears receptive to a wide range of hydrophobic residues (Met, Leu, Ile or Val) typically found at this position in diverse HIV-1 strains. By contrast, direct interactions involving the light chain are limited to Asp50 projecting from CDR L2, which makes a hydrogen bond with Arg500 (Lys446 in HXB2), a residue abutting the Furin recognition site within the C-terminal region of gp120 and conserved as positively charged (Arg or Lys) in >85% of HIV-1 stains.

**Figure 2.**
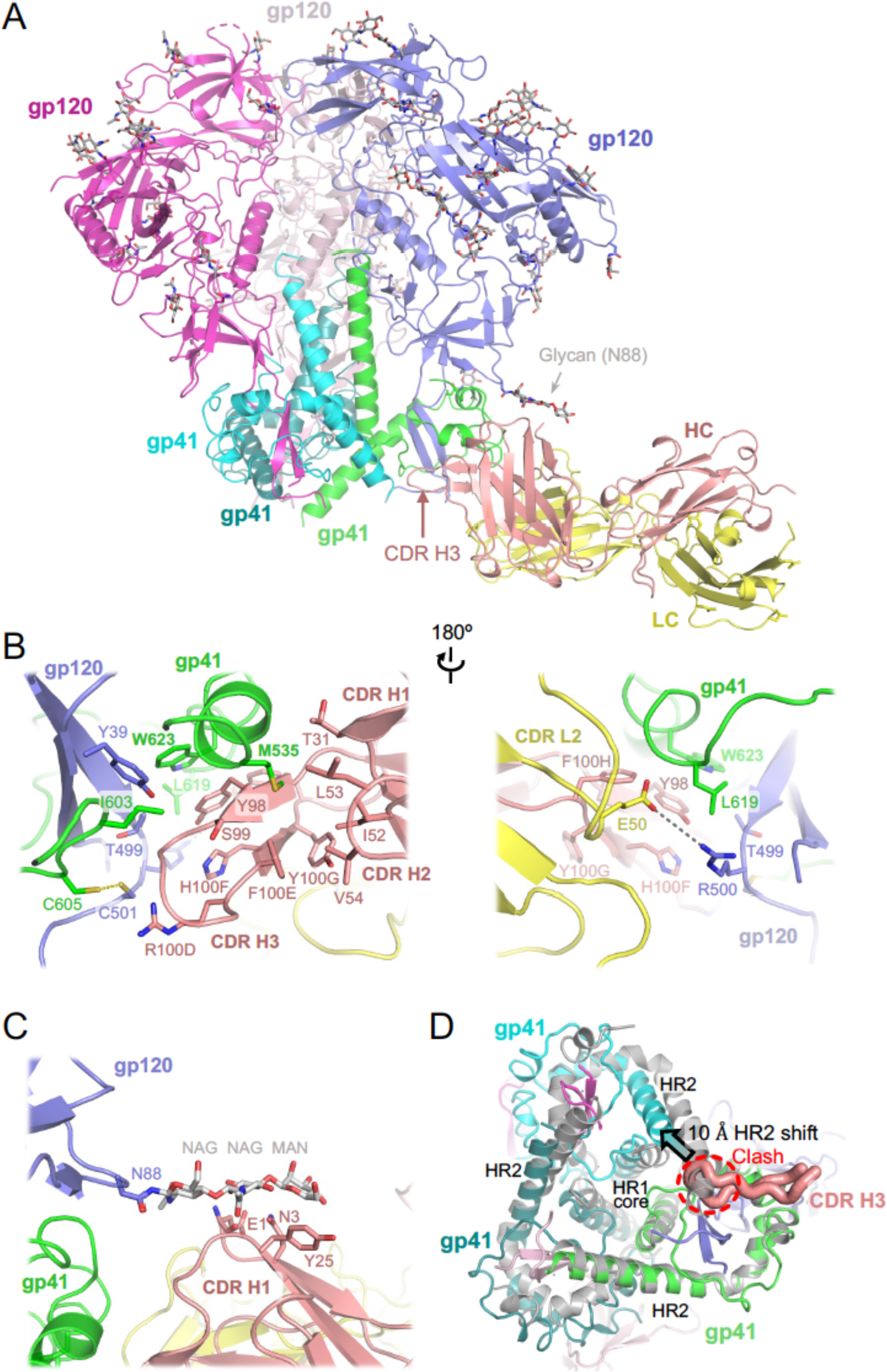
Cryo-EM structure of HIV-1 Env BG505 SOSIP.664 construct in complex with ELC07 Fab. (A) Overview of the atomistic model. The protein chains are shown as cartoons, with individual subunits indicated and colour-coded: gp41 in green, cyan, and teal; gp120 in blue, light pink, and bright magenta; ELC07 heavy chain (HC) and light chain (LC) in salmon and yellow, respectively. The glycans are shown as sticks with carbon atoms in grey. Arrowheads indicate the glycan attached to Asn88 of the gp120 subunit interacting with the antibody and the CDR H3 loop of the heavy chain. (B) A zoomed view on the Env-ELC07 interface shown in two orientations related by a 180° rotation. Side chains of residues discussed in the text are indicated and shown as sticks. ELC07 heavy chain residue numbering follows the Kabat conventions. Yellow dash indicates the designed SOS bond between gp120 Cys501 and gp41 Cys605; grey dash indicates a salt bridge between gp120 Arg500 and ELC light chain Glu50. (C) Zoomed view on Ans88-linked glycan and its interface with ELC07. (D) Structural changes in the HIV-1 Env induced by ELC07 binding revealed by superposition with 3-fold symmetric BG505 SOSIP.664 (PDB ID 4TVP, shown as grey cartoons) ^47^; viewed from the base of the Env ectodomain. CDR H3 is shown as thick ribbon and indicated, with the rest of the Fab structure hidden for clarity. The heptad repeats 1 and 2 (HR1 and HR2) of gp41 subunits are indicated. Insertion of CDR H3 loop is made possible by ∼10 Å shift (indicated by arrowhead) of HR2 belonging to the gp41 chain engaged by the antibody. The position of HR2 complying with the expected 3-fold symmetry is not possible due to a clash with CD HR3 (indicated).

We note that ELC07 binding induces considerable perturbation in the gp41 trimer structure. Directly engaging one of the gp41 chains, CDR H3 inserts into the space occupied by the neighboring gp41 subunit, displacing the HR2 helix by ∼10 Å from its normal position and causing a disorder of eight HR2 C-terminal residues (Fig 2D). The conformational rearrangements propagate throughout the trimer (Fig 2D), likely affecting the stability and function of the viral glycoprotein. The ELC07 epitope is distinct from two well-characterised bnAbs targeting the gp41-gp120 interface, 8ANC195 and 35O22, that show no and partial overlap with ELC07, respectively (Fig S2C). By contrast, the ELC07 binding site appears to more substantially overlap with the epitope of 3BC315, a bnAb that predominantly targets gp41^45^ (Fig S2C) and displays competition in our HIV-1 Env binding assays (Fig S1). Although the interactions of 3BC315 with HIV-1 Env have not been described in atomistic detail, the activities of both ELC07 and 3BC315 are impacted by glycosylation of Asn88 (Fig S1D)^45^. The highly conserved glycan is well-ordered in our cryo-EM structure (Fig 2A; Fig S2A, B) making contacts with the Glu1, Asn3 and Tyr25 of the ELC07 heavy chain (Fig 2C). However, as noted previously^45^, in its natural conformation, the Asn88 glycan likely restricts the access to the epitope explaining its negative effect on the activity of both antibodies.

### Memory B cell subsets of the bnAb donor do not show expected HIV-1-associated dysfunction despite HIV-1 viraemia

Our functional and structural studies demonstrated the ability of the T125 donor to produce bnAbs. Next, we proceeded to examine memory B cell phenotypes in this bnAb donor and explore alterations in B cells with different neutralisation capacities. Firstly, the cell surface markers CD27 and CD21 were used to identify memory B cell phenotypes and, with the inclusion of IgG in analyses, to identify class-switched, *bona fide* memory cells. Surprisingly, IgG+ memory B cells within PBMCs of the bnAb donor were predominantly RM (CD27+ CD21+, ∼60%), with TLM (CD27-CD21-) the most infrequent phenotype (Fig 4A, B). This result was consistent across both timepoints (Fig S3), despite the high viral load of 73,300 copies/ml at the first timepoint. This is in marked contrast to the well-established observation that HIV-1 viraemia leads to a decrease in RM (CD27+ CD21+), and an increase in TLM (CD27-CD21-) and AM (CD27+ CD21-)^33^. To further validate the unexpected memory B cell profile in the T125 donor, we investigated a donor with a similar level of HIV-1 viraemia (110,000 copies/mL) but without bnAbs, and an HIV-negative donor. Consistent with previous literature, there was a pronounced increase in the percentage of IgG+ TLMs in the donor with viraemia (22.4%) relative to the HIV-negative donor (4.9%), in contrast to the bnAb donor (5.9%) who also had viraemia (Fig 4A). Furthermore, the percentage of IgG+ B cells with an RM phenotype was reduced in the donor with viraemia (49.9%) compared to both the HIV-negative donor (74.6%) and the bnAb donor (62.8%). Given previous studies of individuals living with HIV-1 viraemia have reported Env-reactive cells are enriched in the TLM subset^37^, we next examined the phenotype of the Env+ IgG+ B cells. Strikingly, no enrichment in Env+TLM B cells was observed in the T125 donor, despite blood viraemia of 73,300 copies/ml. By contrast, 84% and 96% of Env+ B cells had an RM phenotype at the first timepoint and second timepoints, respectively (Fig 3C, D), while less than 1% were identified as TLM at either timepoint (Fig 3C, D, Fig S3). Together these results show that the memory B cell pool in the bnAb donor is unusual in an individual with HIV viraemia and characterised by an enrichment of HIV-specific RM cells.

**Figure 3:**
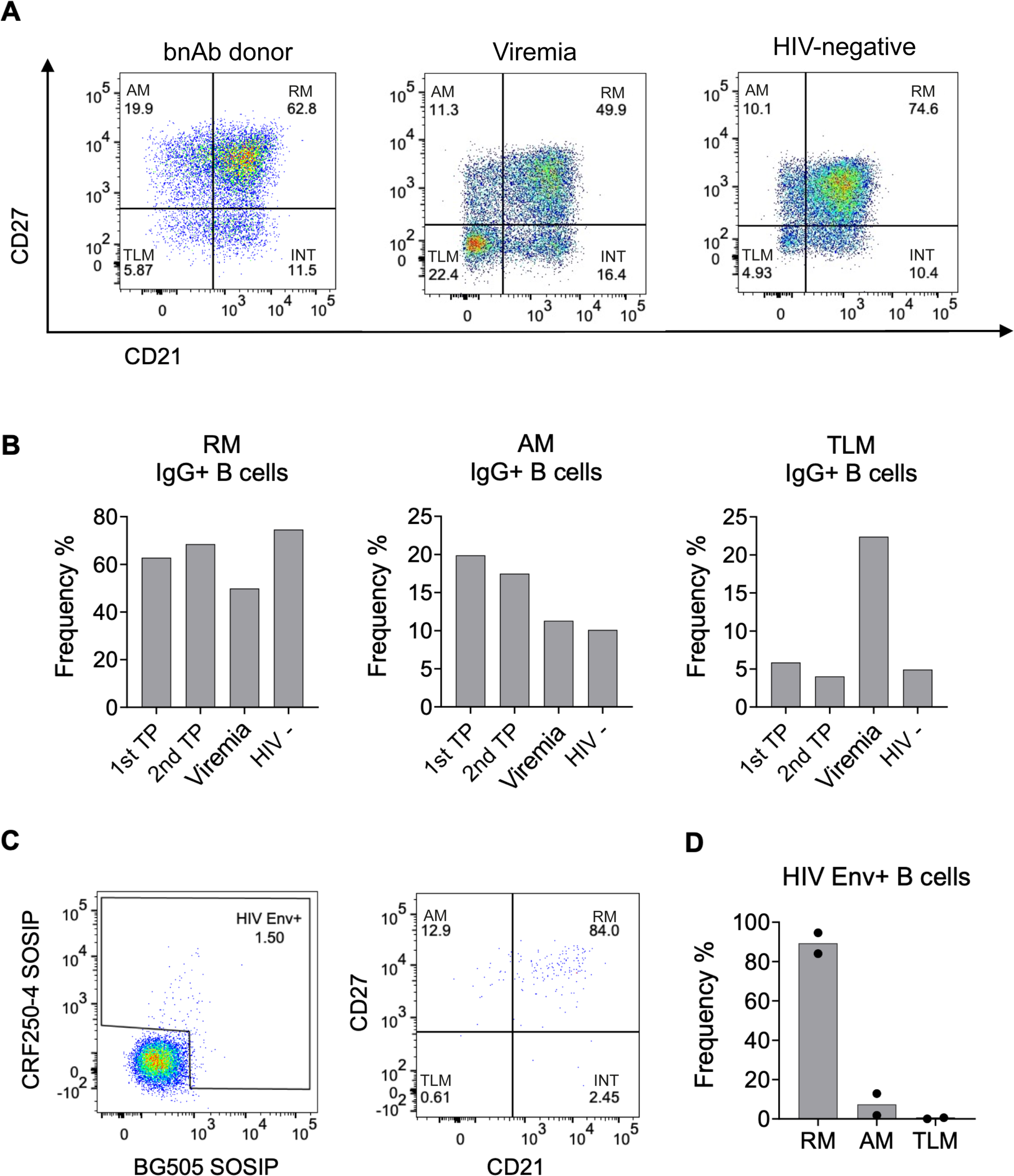
B Cell surface profiles of bnAb donor do not show expected HIV-1-associated dysfunction. (A) FACS analysis of CD27 and CD21 on IgG+ B cells (CD19+ IgG+ IgM-) from PBMC isolated from the bnAb donor T125 (1st timepoint (TP)), an individual living with HIV-1 with detectable viraemia and an HIV-negative donor. (B) Percentage frequency of resting memory (RM), activated memory (AM) and tissue-like memory (TLM) IgG+ B cells in the bnAb donor T125 1st TP, second TP four months later (2nd TP), donor with detectable HIV-1 Viremia and HIV-negative donor PBMC. (C) Percentage of HIV-1 Env+ IgG+ B cells from the bnAb donor T125 1st TP PBMC identified by flow cytometry based on the ability to bind fluorescently labelled CRF250-4 and/or BG505 SOSIP, followed by analysis of their CD27 and CD21 surface expression. (D) Percentage frequency of resting memory (RM), activated memory (AM) and tissue-like memory (TLM) HIV-1 Env+ IgG+ B cells in the bnAb donor T125, with the mean frequency of both TP plotted.

**Figure 4:**
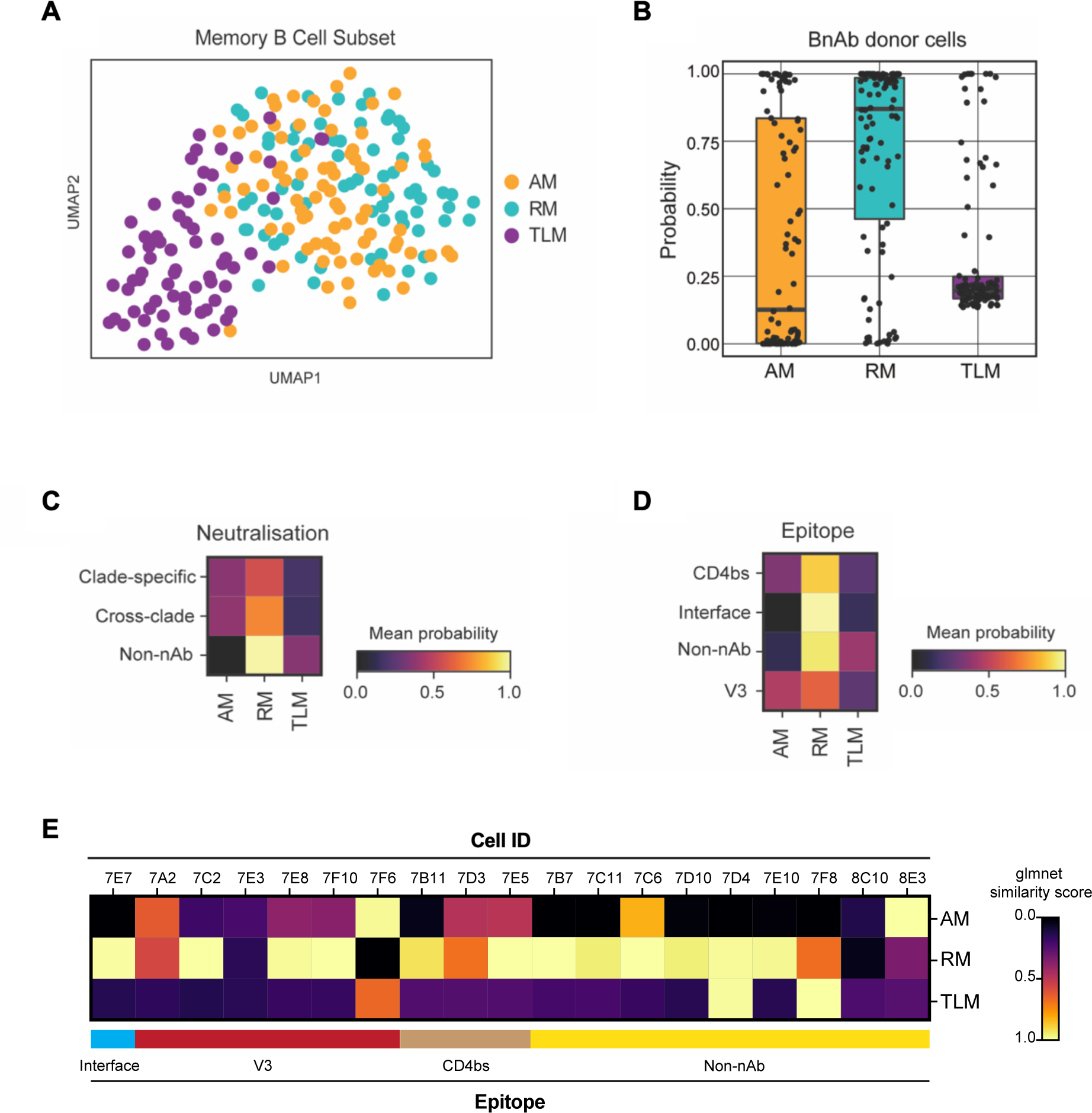
Single B cells from a bnAb donor have a transcriptional phenotype most similar to resting memory cells, irrespective of their BCR specificity or functionality. (A) UMAP visualisation of single-cell transcriptomes (Smart-Seq2) from 223 IgG+ B cells from an individual living with HIV-1 with low VL (100 c/mL at the time of sampling), coloured by their original FACS sorting strategy as resting memory (RM; cyan), activated memory (AM; orange) and tissue-like memory (TLM; purple). (B) Similarity of single-cell transcriptomes of HIV-1 Env reactive IgG+ B cells from the bnAb donor with RM, AM and TLM IgG+ B cell subsets from the low VL donor memory B cell subsets, calculated as a probability using the Glmnet algorithm. (C-D) Heatmaps of the mean probability (as calculated in B) of HIV-1 Env reactive IgG+ B cells from the bnAb donor for memory subsets based on (C) BCR neutralisation of clade C specific or cross-clade HIV-1 PVs or no neutralisation (non-nAb) and (D) BCR epitope targeted on the HIV-1 Env. The BCR specificity and functionality were characterised based on the behavior of soluble mAb cloned and expressed from single B cells from the bnAb donor. (E) Heatmaps of the mean probability (as calculated in B) of each memory subset for each epitope mapped HIV-1 Env reactive IgG+ B cell from the bnAb donor.

### B cells from the bnAb donor have an RM transcriptional phenotype irrespective of their BCR specificity or functionality

To explore whether an RM phenotype was also observed at the transcriptome level and associated with antibody specificity/functionality, we performed Smart-Seq2-based single-cell RNA sequencing of memory B cells from the T125 bnAb donor and an aviraemic individual living with HIV-1 as a reference. Memory B cells were sorted based on expression of CD27 and CD21, allowing their annotation as RM, AM and TLM B cells. Analysis of data generated from the reference donor revealed that TLM B cells were transcriptionally distinct from the RM and AM B cells, as expected (Fig 4A). Next, we trained Glmnet on this reference dataset from the donor with aviraemia and calculated the transcriptomic similarity of each bnAb donor B cell with each B cell in the reference subset. In line with the flow cytometry analysis (Fig 2C, D), we found that the transcriptome of the majority of the bnAb donor memory B cells mirrored RM, with some showing similarity to AM and only very few to TLM B cells (Fig 4B). Moreover, the phenotype with the highest similarity was found to be RM regardless of the neutralisation capacity, epitope, breadth or isotype of the mAbs (Fig 4C, D). Interestingly the one B cell that encoded the bnAb ELC07 (referred to as 7E7 when using cell ID) was found to have the highest probability of being RM (Fig 4E).

To explore the transcriptional differences between B cells from the bnAb donor and the reference donor who was aviraemic, we integrated both single cell datasets. As expected, principal component analysis revealed that the bnAb donor B cells clustered predominantly with RM B cells, with some overlap with AM B cells, but not TLM B cells (Fig S4). Intriguingly, bnAb donor B cells had the highest number of significant differentially expressed genes (DEGs) when compared to TLM B cells, including lower expression of genes previously associated with a TLM phenotype in HIV-1 or atypical B cells in malaria, such as FCRL5, CD19 and ITGAX^51–54^ (Fig S4D). The bnAb donor B cells also had significantly lower expression of genes associated with organisation of secondary lymphoid organs/germinal centers (*LTB*, *CXCR4*) and negative regulation of proliferation (*DUSP1*)^55,56^. Moreover, genes associated with cell proliferation, cytokine expression and BCL-6 suppression (*JUN*, *ZFP36*, *KLF6*, *TXNIP*) were upregulated in both RM and AM B cells^57–59^. Conversely, expression of *MALAT1* (NEAT2) and *IFITM3* were significantly higher in the bnAb donor cells compared to RM and AM B cells, the latter a gene induced by BCR antigen engagement to amplify PI3K signalling^60^. By contrast, only one transcript, *TWIST2*, was significantly more highly expressed in the bnAb donor B cells compared to all three memory B cell subsets, encoding a transcription factor that regulates inflammatory cytokines and induces anti-apoptotic gene expression^61^, potentially underpinning the increased representation of RM cells in this bnAb donor.

### Transcriptomic profiles of single memory B cells from a bnAb donor are distinct from memory B cells from other donors with HIV-1 viraemia

To explore whether the transcriptional differences in bnAb donor memory B cells compared with those in an individual living with HIV-1 without detectable viraemia were directly related to the presence of virus in blood, we integrated two publicly available 10X scRNA-seq PBMC datasets from donors with HIV-1 viraemia (PID529 and PID717), as well as those on ART (PID630 and PID876; both with less than 20 HIV-1 RNA copies/mL)^62^ and 11 control HIV-negative donors (CV0902, 04, 11, 15, 17, 26, 29, 34, 39, 40 and 44)^63^ (Fig S5A). UMAP visualisation revealed that B cells from HIV-negative donors largely formed a separate cluster that only partially overlapped with virally suppressed donors and had the least overlap with donors with HIV-1 viraemia (Fig 5A), indicating that these cells are transcriptionally distinct from those found in HIV-negative controls. Similarly, when considering memory B cells in isolation (Fig S4B-C), cells from people living with HIV-1, whether with suppressed or detectable viraemia, were distinct from the majority of memory B cells from control HIV-negative donors (Fig 5B). Top 10 marker genes analysis of memory B cells from HIV-negative donors included *CD79A*, required for BCR signalling, whereas memory B cells from individuals with detectable viraemia showed genes linked to interferon stimulation, *IFI44L* and I*SG15*, as well as *XAF1*, associated with apoptosis^64^(Fig 5C). B cells from donors living with HIV-1 viraemia had the highest mean expression score of genes associated with hallmark IFN-α response and IFN-γ response (Fig 5D) as expected based on prior reports^62,65^. Similarly, gene set enrichment analysis (GSEA) confirmed a significant enrichment for interferon hallmark genes in this subset (Fig 5E).

**Figure 5:**
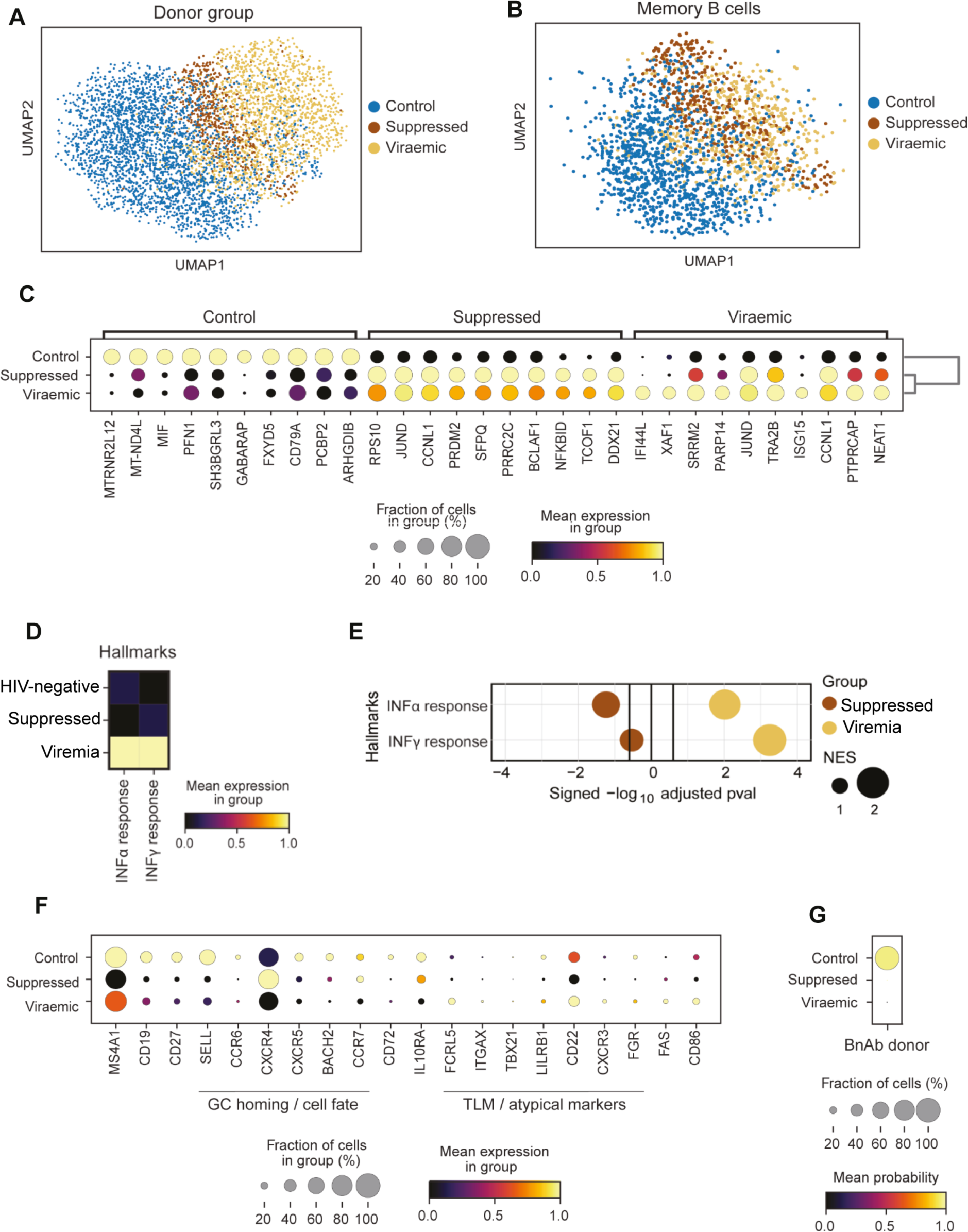
Transcriptomic profiles of single memory B cells from a bnAb donor are unlike memory B cells from other donors with HIV-1 viraemia. (A) UMAP visualisation of B cells integrated from two publicly available scRNA-seq (10x) datasets taken from 11 healthy donors (control) (Stephenson et al. 2021) and 2 donors with detectable viraemia, 2 suppressed donors and 1 healthy donor (Wang et al. 2020), coloured by donor group. (B) UMAP visualisation of memory B cell transcriptomes identified using CellTypist from the single-cell transcriptomes in (A), coloured by their donor group. (C) Expression of top 10 DEGs by memory B cells in each donor group. (D) Hallmark IFN-α and IFN-γ response signature scores of memory B cells from HIV-negative (control), suppressed and donors with detectable viraemia scaled by column. (E) GSEA for a hallmark IFN-α and IFN-γ response based on pre-ranked DEGs in Memory B cells from donors that were HIV-negative (control), suppressed or who had detectable viraemia. Normalised enrichment score (NES) reflects the circle size. Vertical black lines indicate the threshold of significance. (F) Expression of select genes associated with B cell phenotypes in each donor group. The fraction of cells is shown by the dot size and the mean gene expression is reflected by the colour. (G) Dot plot of the CellTypist probability of bnAb donor B cells similarity to memory B cells transcriptomes isolated from donors who were HIV-negative, suppressed or who had detectable viraemia.

Further analysis of selected genes associated with specific memory B cell phenotypes, revealed HIV-negative controls had relatively higher expression of the memory marker CD27 than the people living with HIV-1, whether the donors were suppressed or had detectable viraemia (Fig 5F), consistent with HIV-negative individuals having a higher proportion of RM (CD27^+/high^) B cells. Similarly, there was high expression of *SELL* in memory B cells from HIV-negative donors, which facilitates entry into secondary lymph nodes and *BACH2*, which is required for GC regulation^66^ (Fig 5F). Moreover, expression of the chemokine receptor *CCR7*, linked to GC retention^67^ and in enabling memory B cells to support affinity maturation in the context of antigenic drift^68^, was higher in memory B cells from HIV-negative and virally suppressed donors as compared to memory B cells from donors with viraemia (Fig 5F). Concordantly, people living with HIV-1 who have detectable viraemia have been shown to have higher proportions of TLM cells with low CCR7 expression^52,53^. Activation markers associated with HIV-1 viraemia, namely FAS and CD86^65,69^, were also expressed in a small fraction of the memory B cells from donors with detectable viraemia, but not in those from suppressed or HIV-negative donors (Fig 5F). Based on these profound transcriptomic differences between memory B cells from donors with viraemia, those who were virally supressed, and HIV-negative donors, we assessed their similarity to the bnAb donor B cells. We trained CellTypist on memory B-cell transcriptomes from the three types of donors and calculated the probability (similarity) score for every bnAb donor B cell (Fig 5G). This analysis revealed that memory B cells isolated from the bnAb donor were most similar to memory B cells from HIV-negative control donors, and least similar to memory B cells from people living with HIV-1, whether virally suppressed or those who had detectable viraemia (Fig 5G). Overall, our findings showed at both the transcriptome and proteome level that the memory B cells of this bnAb donor are most similar to RM B cells and lack HIV-1-viraemia associated changes in B cell phenotype, which have been previously suggested to limit functionality^53^.

## Discussion

In this study, we investigated a historical cohort of people living with HIV-1 prior to ART and identified one individual, T125, with highly robust and broad plasma HIV-1 neutralisation. A combination of RNAseq and single B cell cloning allowed us to isolate a bnAb, ELC07, from this donor that displayed ∼50% neutralisation of multiple standard PV panels. We note that in isolation or in combination with the isolated strain specific nAbs, ELC07 did not fully recapitulate its donor’s plasma neutralisation breadth. Therefore, this donor likely produced additional bnAbs that contributed to the elite serum neutralisation, as previously observed in other donors^15,70–73^.

Our cryo-EM study revealed that ELC07 engages a conformation epitope targeting the gp120-gp41 interface within the viral Env glycoprotein trimer. The resulting high-resolution structure provided insight into the broad HIV-1 recognition by ELC07 that have implications for study of other interface bnAbs and their induction by vaccination. Although the structure does not directly explain the mechanism of HIV-1 neutralisation by ELC07 bnAb, it invites several hypotheses for future studies. Firstly, our results demonstrate that ELC07 binding induces considerable perturbations within the gp41 homotrimer at the base of the viral Env, which may cause destabilisation of Env on HIV-1 virions. Indeed, another bnAb with an overlapping epitope, 3BC315, was shown to cause dissociation of HIV-1 Env trimers^45^. Conversely, co-engagement of both gp41 and gp120 subunits by the antibody may restrict conformational rearrangements at the gp41-gp120 interface involved in the opening of the Env structure prior to activation of the fusion machinery^74–77^. Finally, binding at the very base of HIV-1 Env, ELC07 is expected to come in close contact with the viral membrane (Fig S2D). Therefore, the antibody may induce tilting of the viral glycoprotein spike and affect its interactions with the lipid bilayer, as has been proposed with bnAbs targeting the membrane proximal external region of gp41^78,79^.

Unexpectedly, B cell phenotyping revealed that the bnAb donor who was ART naïve and had detectable viraemia did not have a high percentage of total CD27-/CD21-TLM B cells typically found in people living with HIV-1 who are not on suppressive ART^33,65^. Moreover, the vast majority of the donor’s HIV-1 Env-specific IgG+ B cells were found to express both CD27 and CD21, characteristic of RM, a population that is normally markedly reduced in HIV-1 viraemia^52^. Furthermore, single memory B cell transcriptomes from this bnAb donor were most similar to those from HIV-negative donors, with a reduced IFNα and IFNγ response gene set signature relative to people living with HIV-1, whether they had detectable viraemia or were virally supressed. Together, these observations suggest that the preservation of RM B cells and their presumed ability to better respond and mature may allow the development of bnAbs despite the ongoing high antigenic burden of HIV-1 viraemia, which typically leads to B cell dysregulation. Conversely, pharmacological interventions aimed at preservation or restoration of the RM B cell population may benefit the development of bnAbs in people living with HIV-1.

Presumably, the broad and potent plasma neutralisation of donor T125 comprises multiple diverse bnAbs rather than just variants of ELC07, in line with prior studies^15,72,80^. Moreover, the elite plasma neutralisation status of the donor is consistent with existing data linking higher viral load, diversity and time since infection to neutralisation breadth^10,12,21^. However, that neutralisation breadth co-exists with uncontrolled viraemia is not itself intuitively in line with observations of widespread disruption of the memory B cell population during HIV-1 infection ^37,52,81^. Indeed, the current assumption is that individuals make bnAbs despite the widespread B cell dysfunction induced by untreated HIV-1 infection, in the face of evidence that distorted B cell populations are associated with poor antibody responses against other pathogens^82,83^ during HIV-1 infection. Our findings suggest that preserved memory B cell homeostasis may support bnAb development, which agrees with prior data on other facets of the adaptive immune system during bnAb generation. Specifically, higher numbers of circulating T_FH_ cells have been associated with breadth^84^, which also suggests a greater level of immune system preservation in individuals with viraemia who make bnAbs. Indeed, HIV-1 decimates and exhausts CD4+ T cells^85,86^ with a preference for T_FH_, resulting in reduced T cell help to B cells^22^. Furthermore, a gene encoding an endosomal recycling protein RAB11FIP5 is upregulated in NK cells, preventing them from limiting T_FH_ to indirectly support bnAb development^26^. These studies are in line with the model proposed herein, that bnAbs can preferentially develop in a relatively undisrupted immune system with concurrent continual antigenic stimulation due to ongoing viraemia.

Importantly, we have described one particular case of a bnAb donor, who has detectable viraemia and minimal HIV-1-associated memory B cell disturbance. We cannot presume this is required in all cases of bnAb development or would be observed consistently across the complex multi-year development of bnAbs. For example, while viraemia is frequently associated to neutralisation breath, there are exceptions where individuals with low or no viral load display broad neutralisation and can produce bnAbs^87–89^, including the VRC01 donor who was a slow progressor^90^. However, this type of viraemic control could be another scenario, whereby sufficient antigenic stimulation is provided but only locally in secondary lymphoid organs, avoiding widespread peripheral immune disruption, given the reports of ongoing productive viral replication within B cell follicles in individuals living with HIV-1 who are aviraemic^91^.

Our data demonstrate that most Env+ memory B cells in this bnAb donor have an RM phenotype, which contrasts with a previous study showing that IgG+ Env+ (gp140) B cells from 42 people living with HIV-1 who had detectable viraemia were mostly activated (48.8%), with lower levels of RM (37%) and elevated TLM (11.6%)^52^. Notably, the individuals in the cited study also had characteristic global memory B cell disturbance and were not reported to produce bnAbs^52^. More recently, two studies have explored the antibody repertoire in people living with HIV-1 and showed a correlation between higher SHM and neutralisation breadth^30,31^. These studies showed lower average antibody SHM in the HIV-1 group as compared to the HIV-negative group, yet those individuals living with HIV-1 with neutralisation breadth had repertoires that were not perturbed and instead capable of exhibiting similar SHM to HIV-negative participants^30^. This study also revealed a significant negative correlation with the frequency of CTLA-4+ Treg cells and neutralisation breadth, which aligns with prior work associating preserved immune function, in this case higher T_FH,_ with breadth^30^.

Overall, our approach of combining single B cell cloning with a plate-based RNAseq method enabled us to conclude that the bnAb donor B cells exhibited a RM phenotype most similar to HIV-negative donors. While only one bnAb was identified, a range of cross-clade and non-neutralising mAbs were found and there was no substantial variation with B cell phenotype in line with functional activity of the cloned mAbs. Indeed, as most of the Env+ memory B cells had the same RM cell surface phenotype, this implies that there is not a difference between bnAb and non-bnAb B cells, but rather a difference between bnAb and non-bnAb donors. Consequently, we can conclude that the key difference between cells that produced bnAbs and those that did not is the epitope they recognised and not their phenotypic trajectory. This suggests greater GC activity in the bnAb donor than other individuals with viraemia given TLM B cells express homing receptors to inflammatory sites rather than the GC^36^ and previously studied BCRs from Env+ TLM B cells display lower SHM than RM B cells^37^.

Importantly, while the bnAb donor studied here had a strikingly different global B cell profile from that anticipated during HIV-1 viraemia, this is a singular observation to date, albeit is consistent across the two timepoints at which PBMCs were collected. It is unclear whether this individual is an exception or reflective of a more widely shared phenotype that occurs during bnAb development warranting further studies. Given previous observations of greater immune function, in the form of larger T_FH_^84^ and CTLA4+ Treg^30^ populations in other bnAb donors, it seems congruent that preservation of classical RM B cells would also be advantageous for generating bnAbs. However, it has been noted that lower early total B cell numbers are found in those that go on to develop a level of breadth, though these individuals also had greater numbers of founder Env specific B cells^32^. In light of these previous findings, our results suggest that preservation of RM B cells in untreated infection may be advantageous at a particular stage in the generation of bnAbs rather than a prerequisite throughout infection. Unfortunately, no further PBMC samples from this bnAb donor were available to allow greater scrutiny of total (non-Env reactive) memory B cells, non-memory B cells and other important immune cells such as T cells that could have been achieved by 10X genomics, which would ideally be conducted across the course of infection and the development of bnAbs. Future studies will aim to determine the prevalence of the preserved memory B cell homeostasis observed here across other bnAb donors and crucially to understand at which time point during bnAb development limiting HIV-1-associated B cell disturbance is beneficial. However, these results do enable us to conclude that bnAbs can arise without HIV-1-induced dysregulation of the memory B cell compartment and suggest that sufficient levels of antigenic stimulation, appropriately presented to trigger bnAb precursors, should be effective in HIV-negative vaccine recipients.

## Acknowledgments

We would like to thank the participants for their contributions to this study, and the hospital and NHS staff for their assistance in recruitment of patients and acquisition of samples. Sequencing and quality control analysis was performed by the UCL Pathogen Genomics Unit PGU. The authors would also like to thank Marit van Gils for the provision of recombinant Env expressing plasmids and critical reading of the manuscript.

## Author contributions

L.E.M conceptualised the project. S.G, L.M, J.H, C.P, J.M.G, A.F, E.T, C.R-S, A.N, C.R, Y.A, D.F, K.J.D, L.E.M, P.C performed experimental work. S.G, L.M, O.S, Z.K.T, M.C designed and performed bioinformatic analysis. C.O, J.D, J.A, R.K.G, A.M recruited participants. S.G, L.M, O.S, P.C and L.E.M wrote the original manuscript. S.G, L.M, J.H, J.M.G, C.R-S, D.F, K.J.D, C.O, P.C, A.M, M.C and L.E.M reviewed and edited the manuscript. L.E.M, A.M and P.C acquired funding and supervised the project.

## Materials & methods

### Study samples

The protocol for the East London Cohort study was approved by local research ethic committee (06/Q0603/59) for participants recruited at the Grahame Hayton Unit, Barts and The London Hospital; Centre for the Study of Sexual Health and HIV, Homerton University Hospital; and Andrewes Unit at St Bartholomew’s Hospital. Additional samples from people living with HIV-1 who had detectable viraemia were collected as part of a protocol approved by local research ethic committee (London – City & East REC 12/LO/1572). Samples from people living with HIV-1 on ART (suppressed) and HIV-negative participant samples were previously collected and processed with ethical approval by South Central – Hampshire B (REC 19/SC/0423)^83^. The study complied with all relevant ethical regulations for work with human participants and conformed to the Helsinki declaration principles and Good Clinical Practice (GCP) guidelines. All subjects enrolled into the study provided written informed consent. Plasma and PBMCs from the East London cohort were previously collected and cryopreserved with ethical approval (06/Q0603/59)^39^. All participants in the East London cohort had acquired HIV-1 a minimum of one year prior to sampling and were recruited before 2010, meaning that ART was only initiated for those with an AIDS-defining illness or a persisting CD4 cell count <200 cells/μL^92^. Previously generated ID_50_ values against various PV in the TZM-bl assay^39^ were reviewed and samples that neutralised more than one tier 2/3 virus with an ID_50_ titer >100 without MLV reactivity were selected for additional screening of neutralisation breadth.

### Neutralisation assays

Serum and plasma samples or monoclonal antibodies (sterile filtered, 0.22 μM) were titrated 2-fold or 3-fold down a 96-well flat-bottom white plate (Thermo) containing complete DMEM (leaving wells without sample for virus and cell only controls) and then incubated with a 200 TCID_50_ dilution of PV for 1 hr at 37°C. Serum and plasma samples were diluted prior to titration to have a starting dilution of 1:100 (or 1:75 or 1:50) after the addition of PV. mAbs were used at different starting concentrations (0.5 μg/mL −10 μg/mL) depending on their potency. HeLa TZM-bl reporter cells (1×10^4^ cells/well) containing 25 μg/mL DEAE dextran were added and incubated for 48 hrs in a 37°C incubator with 5% CO2. Media was removed from each well prior to addition of 100 μL Bright-Glo™ luciferase substrate (Promega) diluted 1:20 in 1x lysis buffer. The luciferase activity in RLU was measured using a PheraStar Plus microplate reader (BMG Labtech). Serum and plasma 50% inhibitory dilution (ID_50_) values were calculated from sigmoidal dose-response curves using GraphPad Prism software. Neutralisation scores were calculated from log-transformed titers as in^40^, using the equation Y = log3 (dilution/100) + 1.

### Epitope mapping

To detect changes in neutralisation potency single point mutations were introduced into *env* encoding plasmids for PV production using the QuikChange Lightning Site-Directed Mutagenesis (SDM) kit (Agilent) according to the manufacturer’s protocol. For neutralisation absorption assays commercially produced MPER peptide (ELLELDKWAS LWNWFGITKWLWYIKIFIM, synthesised by Smart BioScience) or in house produced recombinant gp120 D368R was added to serum/mAbs prior to the addition of PV in the neutralisation assay. For competition ELISA, unlabelled mAbs were pre-incubated with blocked lectin immobilised Env. Biotinylated mAbs were then added for 1h, followed by streptavidin-AP detection.

### Protein production

HEK-293F cells (1×10^6^ cells/mL in an Erlenmeyer flask) were transfected with mAb, SOSIP or gp120 encoding plasmids. PEI-MAX was added to sterile filtered (0.22 μM) plasmids and OptiMeM, then left to incubate for 20 mins at room temperature before transfection. For in vivo biotinylation of SOSIP with an avi-tag, 8 mL of the transfection mix was added to HEK-293F cells (200 mL) along with 3 mL of 10 mM biotin. Env proteins were purified by lectin affinity chromatography followed by size-exclusion chromatography (Superdex 200 Increase 10/300 GL column). mAbs were purified by affinity chromatography using protein G resin.

### Cryo-EM sample preparation, data collection and structure refinement

A DNA fragment encoding ELC07 Fab heavy chain with a C-terminal hexahistidine (His_6_) tag and subcloned into pcDNA3.1 was generated by GenScript. The constructs used for expression of stabilised trimeric HIV-1 Env (BG505 SOSIP.664) and Furin have been described^49^. HIV-1 Env and ELC07 Fab used for cryo-EM were produced by transient transfection of Expi293 cells with endotoxin-free preparations of recombinant plasmids using ExpiFectamine 293 (Fisher Scientific). To produce ELC07 Fab, Expi293 cells (400-ml culture grown to a density of 3.5×10^6^ cells per ml) were co-transfected with the plasmids encoding non-tagged light chain and His_6_-tagged Fab heavy chain (used at a molar ratio of 1:1). Secreted recombinant protein was purified from conditioned medium 5-days post-transfection by affinity capture on Ni-Sepharose Excel resin (Cytiva Life Sciences). Following extensive washing, His_6_-tagged Fab was eluted with 200 mM imidazole in 250 mM NaCl, 25 mM Tris-HCl, pH 7.4. The protein was further purified by size exclusion chromatography through a Superdex-200 column (Cytiva Life Sciences) in PBS. To produce BG505 HIV-1 Env SOSIP.664, Expi293 cells (1-L culture grown to a density of 3.5×10^6^ cells per ml) were co-transfected with plasmids expressing BG505 SOSIP.664 and Furin (used at a ratio of 4:1). Five days post-transfection, the protein secreted into conditioned medium was captured onto *Galanthus nivalis* lectin agarose (Vector Laboratories). Following extensive washes with 0.5 M NaCl in PBS, the protein was eluted with 1 M methyl-alpha-D-mannopyranoside in PBS. Trimeric BG505 SOSIP.664 was further purified by size exclusion chromatography through a Superdex 200 column equilibrated with 150 mM NaCl, 50 mM Tris-HCl, pH 7.5.

For vitrification, 4 μl BG505 SOSIP.661 homotrimer at a final concentration of 0.55 mg/ml, supplemented with 0.48 mg/ml ECL07 Fab and 0.085 mM n-dodecyl β-D-maltoside was applied to glow-discharged 400-mesh R1.2/1.3 C-flat holey carbon grids (Electron Microscopy Sciences; product code CF413-50-Au) for 1 min, under 100% humidity at 20°C, before blotting and plunge-freezing in liquid ethane using Vitrobot Mark IV (Thermo Fisher Scientific). Cryo-EM data were acquired on a 300-kV Titan Krios G2 cryo-electron microscope equipped with a Falcon 4i direct electron detector and a Selectris energy filter (Thermo Fischer Scientific). Micrographs were recorded in dose-fractionation mode, at a calibrated magnification corresponding to 0.95 Å per physical pixel. 1,674 EER frames collected per micrograph movie were subsequently processed in 54 fractions, with an exposure dose of 1.06 e/Å^2^ per fraction. A total of 31,737 micrograph movies were collected using an energy filter slit width of 10 eV and a defocus range set at −1.3 to −3.1 µm.

The movie stacks were aligned and summed, with dose weighting, as implemented in Relion-4.0^93,94^). Contrast transfer function (CTF) parameters were estimated using Gctf-v1.18^95^. At this stage, micrographs with crystalline ice contamination were discarded, and the remaining 31,602 images were retained for further processing. An initial set of particles picked with SPHIRE-crYOLO using general model^96^ was subjected to reference-free 2D classification in cryoSPARC-4.3^97^. Particles belonging to well-defined 2D classes were used to train a model for particle picking in Topaz^98^. Picking the entire set of micrographs using Topaz resulted in 4,270,949 particles, which were extracted with a pixel size 3.8 Å and a box size of 90 pixels and subjected to iterative rounds of 2D classification in cryoSPARC-4.3. 877,871 particles contributing to well-defined 2D class averages were re-extracted with a pixel size of 1.9 Å and subjected to 3D classification in Relion-4.0 into five classes using an initial model generated using *Ab-initio* reconstruction in cryoSPARC-4.3. The classification revealed two well-defined 3D classes representing trimeric HIV-1 Env ectodomain with a single Fab molecule bound. 407,874 particles contributing to these classes were re-extracted with the original micrograph pixel size of 0.95 Å and a box size of 360 pixels and used for *Ab-initio* reconstruction in cryoSPARC-4.3 into six 3D classes. 389,761 particles representing well-defined 3D classes were used for Non-uniform refinement in cryoSPARC-4.3 followed by 3D classification without realignment in cryoSPARC-4.3 (into ten 3D classes) and *Ab-initio* reconstruction (four classes) resulting in the final set of 275,291 particles. The final 3D reconstruction (Fig S2B) was obtained by Non-uniform refinement in cryoSPARC-4.3 following Bayesian particle polishing as implemented in Relion-4.0 and a local CTF refinement in cryoSPARC-4.3. The resolution metrics reported here are according to the gold-standard Fourier shell correlation (FSC) 0.143 criterion^99,100^ (Fig S2A). Local resolution of the 3D reconstruction was estimated in cryoSPARC-4.3 (Fig S2A). For illustration purposes and to aid in model building, the cryo-EM map was processed with DeepEMhancer using the tight target model^101^; for real-space refinement of the atomistic model, the reconstruction was sharpened and filtered as implemented in cryoSPARC-4.3 based on local resolution metrics.

The initial atomistic model was generated using HIV-1 Env structure from PDB entry 8FR6 ^102^ and ELC07 Fab model produced by AlphaFold2 Multimer version 3^103^ via ColabFold ^104^. The antibody residues were numbered following Kabat conventions using Abnum tool (http://www.bioinf.org.uk/abs/abnum/)^105^. UCSF Chimera^106^ was used for the initial rigid body docking. All glycan residues were removed and the model was subjected to further refinement using six rigid bodies (one per each Gp120 and Gp41 protein chain) in phenix.refine version 1.21rc1-5084^107^ followed by flexible fitting in Namdinator (https://namdinator.au.dk)^108^ using default parameters. The model was improved by iterative manual building and real-space refinement in Coot^109^. Asn-linked glycan residues were added when supported by the cryo-EM map. The artificial intersubunit SOS disulfide bond between Cys residues 501 and 605 within the SOSIP.664 construct was not unambiguously supported by the cryo-EM maps, and the residues remained unlinked in the model. The final model, refined in real-space using phenix.refine version 1.21rc1-5084, had good fit to the cryo-EM map and reasonable geometry as assessed by MolProbity^110^ (http://molprobity.biochem.duke.edu) (Table S1). Locally filtered cryo-EM maps along with the original half-maps as well as the final refined model will be deposited with the EM and Protein Data Banks upon provisional acceptance of the manuscript.

### Cell staining and phenotypic analysis

PBMCs were thawed, added to complete DMEM and pelleted by centrifugation at 800g for 5 mins. The cell pellet was washed with PBS, pelleted again (800g for 5 mins) and then cells were counted under a microscope using a haemocytometer. Zombie Aqua dead cell stain (1 µl in 400 µl PBS) was added per 1×10^7^ cells and left to incubate for 20 mins, protected from light. Complete DMEM was added to quench the stain, then cells were pelleted (800g for 5 mins) and washed with PBS before adding 100 µl of antibody cocktail per 5×10^6^ cells as follows: CD4 BV510, CD19 FITC, CD21 PE-Cy7, CD27 BV421, IgM APC-Cy7, IgG APC and incubated for 30 mins at room temperature, in the dark. For antigen-specific cell staining, 3 µg of biotinylated SOSIP Env probes were incubated at room temperature (for 30 mins with streptavidin-conjugated fluorophores prior to adding to the antibodies cocktail above. After staining, cells were washed with PBS and then resuspended in PBS. Stained PBMCs were analysed by flow cytometry using a BD FACS**-**Aria or BD FACS-Melody and data were visualised and gated using FlowJo v10.7.1.

### Isolation of single B cells for mAb cloning and scRNA-seq

Single IgG+ antigen-specific (HIV-1 Env+) B cells were isolated using a BD FACS-Melody, with the purity threshold set to yield. Cells were sorted one per well into a 96-well plate containing lysis buffer (0.2% Triton X-100 and RNase inhibitor), oligo-DT primers and dNTPs. The Smart-Seq2 protocol was followed to carry out full-length scRNA-seq on single memory B cells, with modification of the pre-amplification step that was optimised to reduce primer-dimer by excluding the IS PCR primers from the PCR mix. Briefly, total mRNA in each well was reverse transcribed and then pre-amplified using 18 PCR cycles to generate cDNA that was purified using Ampure XP beads. The purified cDNA was assessed by Agilent Tapestation to confirm a peak at 1-2 kb and was quantified by Qubit for normalisation to 1.5 ng for optimal tagmentation (adjusted from 1 ng to account for the presence of primer-dimer). Libraries for sequencing were then generated by performing an enrichment PCR of 12 cycles using an Illumina Nextera XT DNA Library Preparation kit with 96 indices, then assessed by Agilent Tapestation to confirm a peak at 300-800 bp and quantified by Qubit for normalisation to 5 nM. The pooled libraries were submitted to the UCL Pathogen Genomics Unit for sequencing on Illumina NextSeq 500 with 75 bp paired-end reads. The pre-amplified and purified cDNA was also used as the starting material for nested PCRs (PCR1 and PCR2) to amplify the antibody variable regions from heavy, kappa or lambda sequences from each IgG+ B cell as previously described^111^. Recombination-based cloning was conducted using NEBuilder HiFi Assembly Master mix, according to the manufacturer’s (NEB) protocol to insert an unpurified antibody V-region into human mAb expression vectors as previously described^111,112^.

### scRNA-seq data processing and quality control

The Smart-Seq2 library had a median count depth of 1.3 million reads/cell, with a median of 1370 unique genes/cell in line with previous B cell libraries generated using Smart-Seq2^113^. The Smart-Seq2 sequencing data were mapped to the GRCh38 reference human genome in Ensembl version 84, using the STAR algorithm. The transcript and gene abundance were estimated using RSEM^114^ to generate a count matrix. Data were then analysed by isOutlier to assess the quality of libraries based on the count depth, the number of genes detected and the percentage of mitochondrial genes. BCR sequences were assembled from V(D)J transcripts using BraCeR^115^.

### Smart-Seq2 single-cell data analysis

Data were then processed using scanpy (v.1.9.1) workflow with standard quality control steps; cells were filtered if number of genes >6000 or <600. Mitochondrial content was determined using scanpy.pp.calculate_qc_metrics function; cells with mitochondrial genes percentage <50% were retained for further analyses. Genes were retained if they were expressed by at least 2 cells. Gene counts for each cell were normalised to contain a total count equal to 106 counts per cell. This led to a working dataset of 98 cells from the bnAb donor and 223 cells from the aviraemic donor. The top 2000 highly variable genes were selected based on Seurat v.3 algorithm (flavor = seurat_v3) with batch key “Sequencing_batch”. Highly variable genes were further refined by removing potentially confounding genes using the following search formula: ‘^HLA|^IG[HKL][VDJC]|^MT|^A[A-Z][0-9]|^B[A-Z][0-0]’. The number of principal components used for neighbourhood graph construction and dimensional reduction was set at 20. Data integration from both donors was performed using the bbknn algorithm^116^. Uniform Manifold Approximation and Projection (UMAP; v3.10.0)^117^ was used for dimensional reduction and visualisation with all parameters as per default settings in scanpy. For the assessment of transcriptional similarity between cells from bnAb donor and reference cell subsets, Glmnet^118^ and Celltypist^119^ packages were used. For Glmnet-based probability scores, trainScSimilarity/predScSimilarity functions from kelvinny tools (https://github.com/zktuong/kelvinny) were used with alpha set at 0.9 and nfolds 10. Celltypist models and probability scores were generated as per default settings. Differentially expressed genes between bnAb donor B cells and aviraemic donor B cell subsets were assessed using scanpy.tl.rank_genes_groups function based on Wilcoxon rank sum test.

### Public single-cell datasets processing and analysis

^62^: data from donors HD1, PID471, PID529, PID630, PID717 and PID876 were concatenated using anndata ^120^ and processed with scanpy as described above with the following changes: cells were filtered if number of genes >3000 or <200, mitochondrial genes percentage >30%. Genes were retained if they are expressed by at least 3 cells. Gene counts for each cell were normalised to contain a total count equal to 104 counts per cell. Celltypist (model: Immune_All_Low.pkl) with majority voting was used to identify B cells. Raw B cell data were then exported as a separate h5ad object.

^63^: all HIV-negative donor data from Cambridge were concatenated and processed as described for ^62^ data above.

Raw B cell data objects exported from ^62^ and ^63^ datasets were subsequently concatenated and processed by scanpy QC workflow leading to a working dataset of 4941 B cells. Top 2000 highly variable genes were selected based on Seurat v.3 algorithm (flavor = seurat_v3) with batch key “dataset” and refined by removing the following genes ‘^HLA|^IG[HKL][VDJC]|^MT|^A[A-Z][0-9]|^B[A-Z][0-0]’. Bbknn was used for datasets integration with batch_key = ‘dataset’. Celltypist (model: Immune_All_Low.pkl) with majority voting was used to identify memory B cells. These memory B cells data were then used for training a new Celltypist model (with feature_selection set as TRUE and check_expression as FALSE) allowing label transfer to query B cell data (bnAb HIV-1 donor) as control, viraemia or suppressed, respectively.

Interferon α and γ response score was created by using scanpy.tl.score_genes with the reference gene sets being GSEA Hallmark ‘interferon alpha response’ and ‘interferon gamma response’. Gene set enrichment analysis (GSEA) was performed using the fgsea package available on Bioconductor and visualised with the GOChord function in the GOplot package. Briefly, genes were ranked in the descending order by the Wilcoxon statistic value from the pairwise Wilcoxon rank sum tests (suppressed vs. resting, viraemia vs. resting). All unique leading-edge genes from the ‘interferon alpha response’ and ‘interferon gamma response’ pathways were then subject to a heatmap visualisation.

## Data availability

De-multiplexed sequencing reads have been deposited on the EMBL-EBI Functional Genomics Data ArrayExpress repository with accession E-MTAB-1359. This paper does not report original code or software. All computational methods used have been referenced and are publicly available.

