## Supplementary information for "Preservation of memory B cell homeostasis in an individual producing broadly neutralising antibodies against HIV-1"

**Griffith, Muir, Suchanek et al. Supplementary information**

**Figure S1– Clinical characteristics of bnAb donor T125 and and epitope specificity of ELC07**

**Figure S2 – Figure S2. Cryo-EM structure of HIV-1 Env in complex with ELC07 Fab**

**Figure S3– B cell subset distributions in the ELC07 bnAb donor**

**Figure S4 – Comparison of HIV-1 Env reactive IgG+ B cells from BnAb donor and total memory B cells from the reference donor**

**Figure S5 – Integration of publicly available data from HIV-negative controls and individuals living with HIV-1 viremia**

**Table S1 – Cryo-EM data collection, image processing and model refinement.**

**References**

**A**

| Patient ID | T125 |
|---|---|
| Gender | Male |
| Country of birth | Sri Lanka |
| HIV clade | C |
| EC / LTNP | No |
| Year of sampling | 2009 |
| Age at sampling | 37 |
| Viral load | 73700 |
| CD4 count | Unknown |
| Neutralisation score | 3.19 |

**B**

| Epitope | PV mutation or protein absorption | Fold change in $ID_{50}$ titre |
|---|---|---|
| gp120 | BG505 gp120 absorption | 2.5 |
| High mannose patch | JRCSF N332A/N295A | 1.0 |
| Trimer apex | CRF250-4 N160A/K169T | 8.0 |
| CD4bs | JRCSF D279A | 1.0 |
| | BG505 N276D/N462D | 4.2 |
| Fusion peptide | JRCSF T90A | 1.1 |
| MPER | JRCSF MPER absorption | 0.9 |

**C**

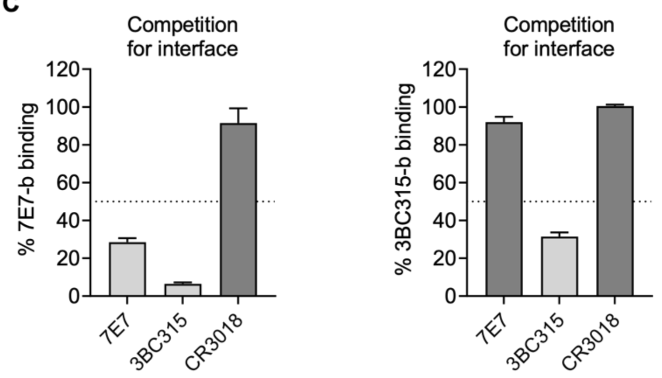


**D**

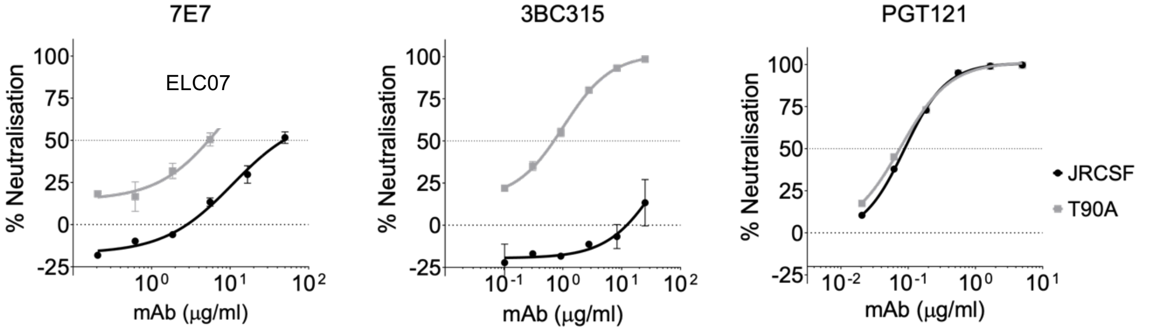


**Figure S1– Clinical characteristics and epitope specificity of bnAb donor T125 and ELC07 – related to Figure 1**

**(A)** Demographic characteristics and clinical features for elite neutraliser T125. **(B)** Effect of protein absorption or PV mutation on plasma neutralisation by elite neutraliser T125, with a 3 fold-change in ID50 titer compared to the wild-type (WT) PV or condition highlighted.

**(C)** Percentage binding of biotinylated ELC07 and 3BC315 to the gp120-gp41 interface of BG505 SOSIP in reciprocal competition ELISAs, mAbs were tested in duplicate with the mean and error bars shown. The non- mAb CR3018 was included as a negative control and a dotted line at 50% indicates the threshold for competitive binding. **(D)** Neutralisation curves of bnAbs ELC07, 3BC315 and PGT121 titrated in duplicate against JRCSF PV (black) and a T90A mutated version (grey) to remove the glycan site at position 88, with the mean and error bars shown. Dotted lines to indicate 0% and 50% neutralisation.

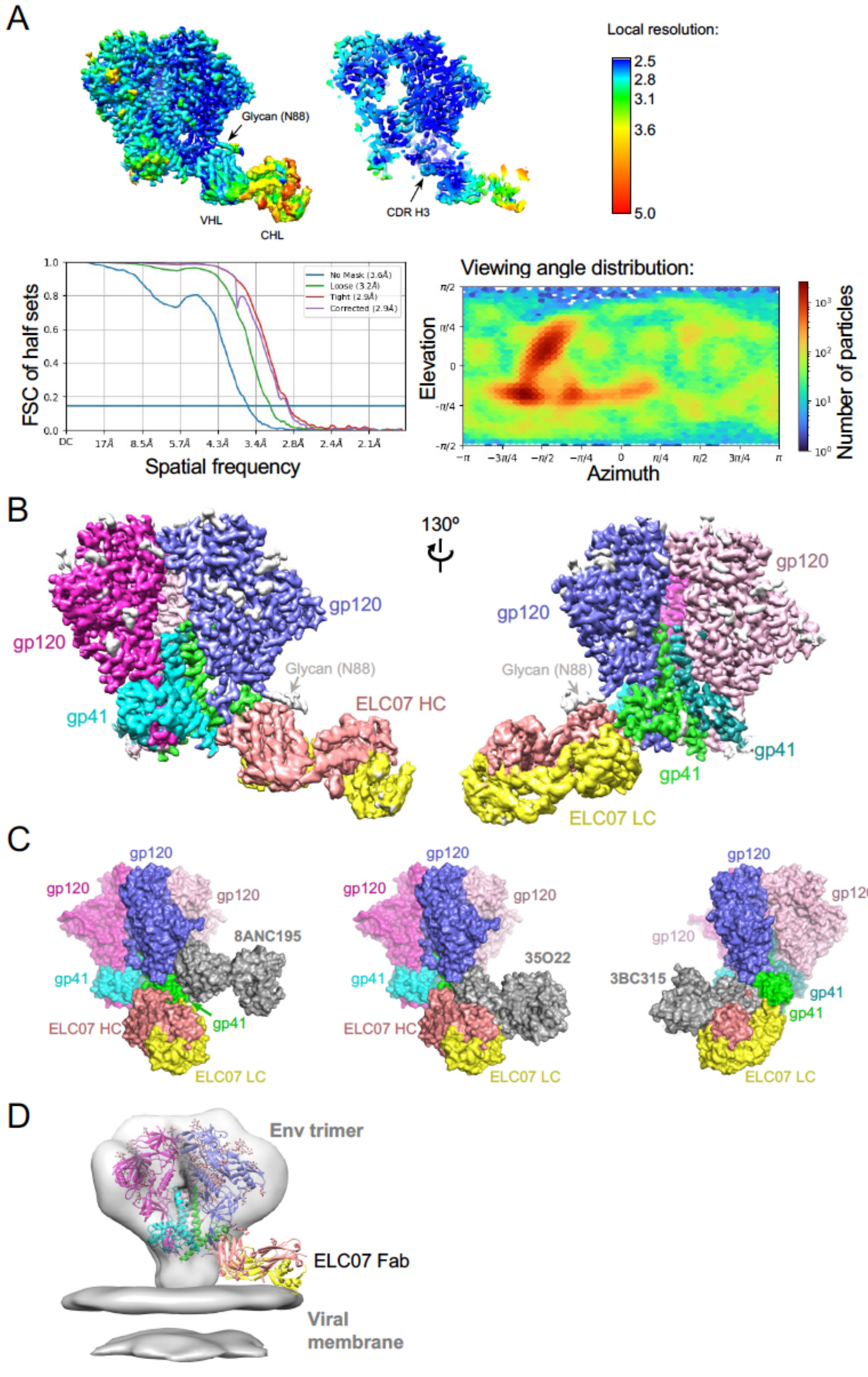


**Figure S2. Cryo-EM structure of HIV-1 Env in complex with ELC07 Fab.**

(A) Top panels show the cryo-EM map locally filtered using DeepEMhancer[1] and coloured by local resolution (as indicated on the inset on the right). The middle panel shows a slice through the map with features representing variable (VHL) and constant (CHL), and the CDR H3 loop indicated. Bottom left panel shows half-map Furrier shell correlation (FSC) without mask (blue line), with loose mask (green line), with tight mask (red line), and optimised mask (purple line). The FCS threshold of 0.143 (horizontal blue line) corresponds 2.9 Å global resolution of 2.9 Å. Bottom right panel shows the viewing angle distribution of assigned to individual particle images that contributed to the final 3D reconstruction.
(B) Cryo-EM map coloured by protein chain as in Fig1, viewed in two orientations related by 130-Å rotation, as indicated.
(C) Comparison of ELC07 to other characterised bnAbs that target the gp120-gp41 interface. The reconstruction of the Env in complex with ELC07 (coloured as in Fig1) is shown superposed on Env in complex with 8ANC195 (PDB ID 5CJX[2], left), 35O22 (PDB ID 4TVP[3], middle), or 3BC315 (EMBD ID 3067, PDB ID 5CCK (26404402), right). Protein chains are shown in space-fill mode with the Env-ELC07 structure coloured as in Fig1; the alternative interface targeting bnAbs in grey.
(D) A model of the ELC07 Fab bound to HIV-1 Env on the viral particle. The Env-ELC07 structure (cartoons, coloured as in Fig1) was docked in the cryo-electron tomography map of HIV-1 spike on in the viral lipid bilayer membrane (EMDB IDs 5019 and 5022 [4], shown as grey surfaces).

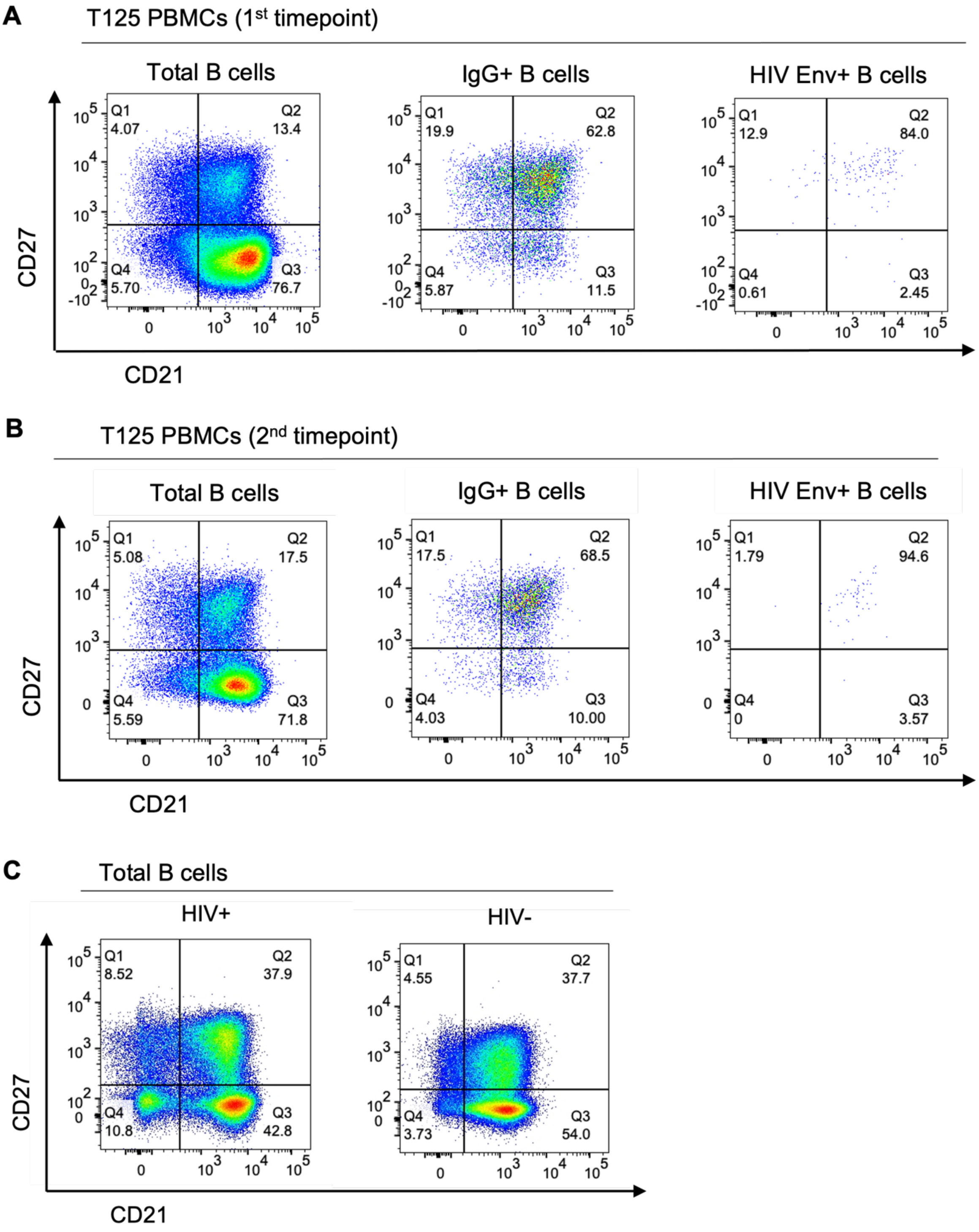


**Figure S3 – B cell subset distributions in the ELC07 bnAb donor – related to Figure 3** **(A-B)** FACS analysis of total B cells (CD19+ CD4-), IgG+ B cells (CD19+ IgG+ IgM-) and Env+ (CD19+ IgG+ SOSIP+) B cells based on the percentage surface expression of CD27 and CD21 from bnAb donor T125 (A) 1st timepoint PBMC and (B) 2nd timepoint PBMC (sampled four months apart). **(C)** FACS analysis of total B cells (CD19+ CD4-) based on the percentage surface expression of CD27 and CD21 from a person living with HIV-1 with viremia and an HIV-1 negative donor.

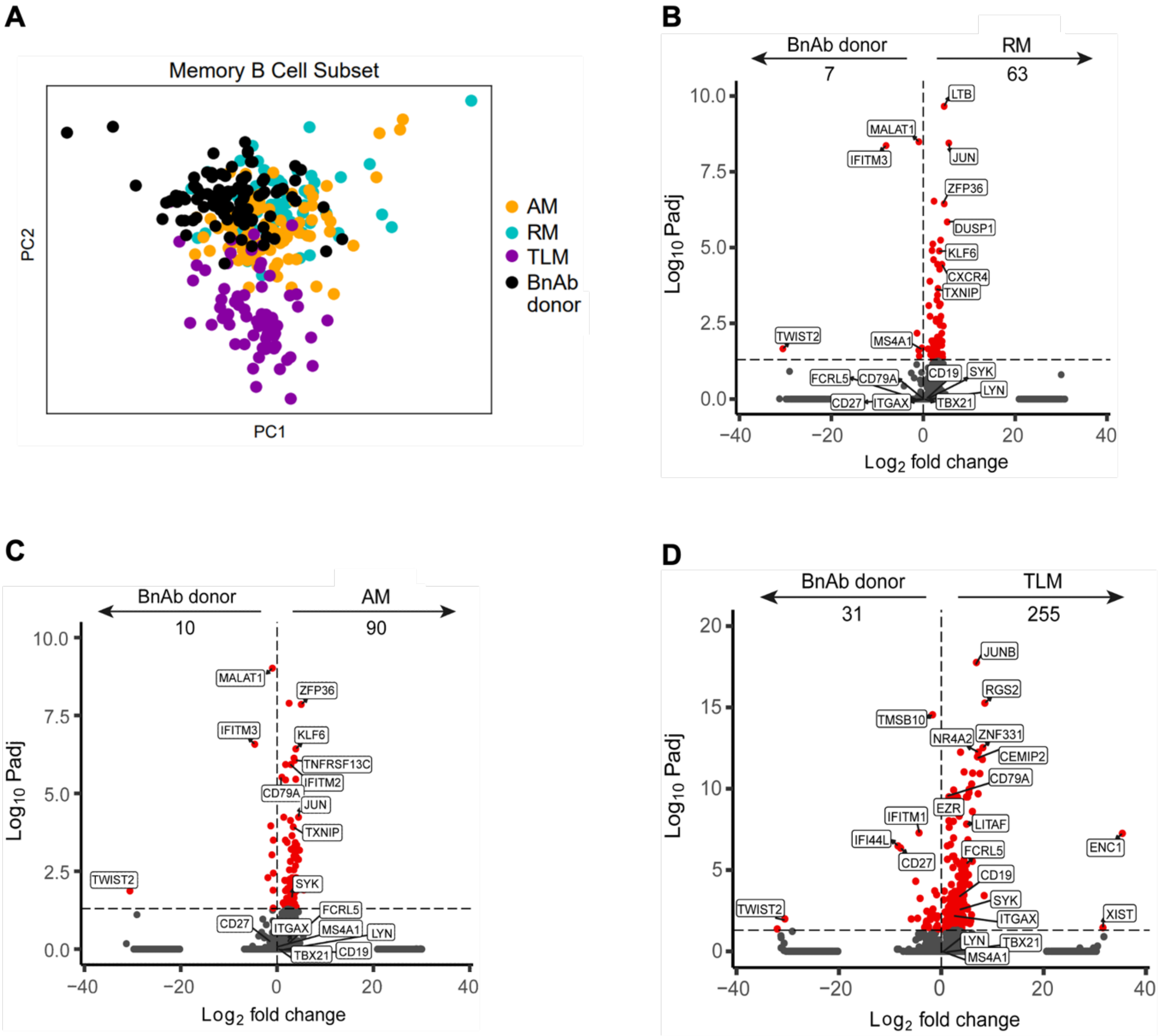


**Figure S4 – Comparison of HIV-1 Env reactive IgG+ B cells from BnAb donor and total memory B cells from the reference donor – related to Figure 4**

(A) PCA visualisation of HIV-1 Env reactive IgG+ B cells from the bnAb donor (black) after integration with resting memory (RM; cyan), activated memory (AM; orange) and tissue-like memory (TLM; purple) IgG+ B cells from a person living with HIV-1 donor with low VL. (B-D) Volcano plots of DEG between HIV-1 Env reactive IgG+ B cells from the bnAb donor and (B) RM (C) AM and (D) TLM IgG+ B cells. The horizontal dotted line indicates the adjusted p-value threshold for significant DEGs that are highlighted in red and the total number of DEGs is stated underneath each arrow. The top DEGs, as well as genes of interest, are annotated.

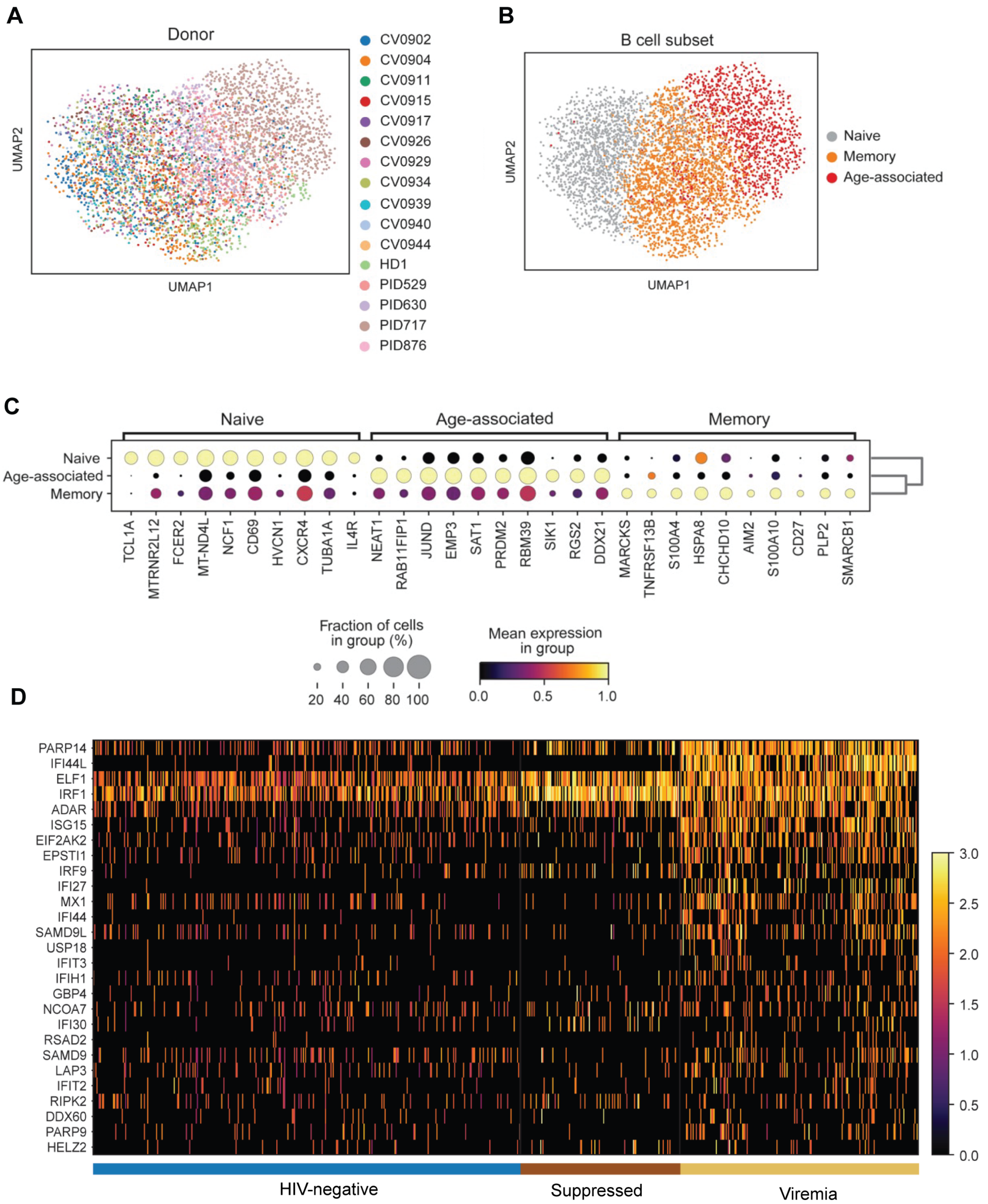


**Figure S5 – Integration of publicly available data from HIV-negative controls and individuals living with HIV-1 viremia– related to Figure 5**

(A-B) UMAP visualisation of B cells integrated from two publicly available scRNA-seq (10x) datasets taken from 11 HIV-negative donors[5] and 2 people living with HIV-1 with viremia, 2 people living with HIV-1 with viral suppression and 1 HIV-negative donor[6]. Coloured by (A) donor ID or (B) Celltypist annotation. (C) Expression of the top 10 DEGs for each B cell subset annotated by Celltypist. The dot size indicates the fraction of cells expressing each particular gene, the dot colour indicates mean gene expression. (D) Expression heatmap of unique leading edge genes from GSEA for hallmark IFN by memory B cells from donors that were HIV-negative (control), suppressed or who had detectable viremia.

**Table S1. Cryo-EM data collection, image processing and model refinement.**

| | |
|---|---|
| **Data collection** | |
| Microscope, operating voltage | Titan Krios G2, 300 keV |
| Detector | Falcon 4i |
| Magnification (nominal) | 130,000 |
| Pixel size (Å) | 0.95 |
| Underfocus range (nominal, μm) | 1.3 - 3.1 |
| Number of EER frames per movie | 1,674 |
| Total electron fluence (e/Å$^2$) | 33 |
| Automation software | EPU |
| Total number of micrograph movies used | 31,602 |
| **Reconstruction** | |
| Software for 2D classification | cryoSPARC-4.3 |
| Software for 3D classification | Relion-4.1, cryoSPARC-4.3 |
| Software for final reconstruction | cryoSPARC-4.3 |
| Number of initially extracted particles | 4,270,949 |
| Number of refined particles | 275,291 |
| Symmetry | *C1* |
| Global resolution (FSC 0.143, Å) | 2.92 |
| Map resolution range (Å) | 2.5 - 4.5 |
| Map sharpening B factor | -115.9 |
| **Model refinement** | |
| Software for real-space refinement | Phenix 1.21rc1-5084 |
| Model composition | |
| Number of non-hydrogen atoms | 17,517 |
| Number of protein residues | 2,113 |
| Number of glycan residues (NAG, BMA, MAN) | 67, 4, 4 |
| *B* factors (Å$^2$) | |
| Protein | 54.1 |
| Glycan residues | 67.4 |
| Real-space correlation coefficient ($CC_{mask}$, $CC_{box}$, $CC_{peaks}$, $CC_{volume}$) | 0.84, 0.73, 0.73, 0.80 |
| R.m.s. deviations | |
| Bond lengths (Å) | 0.004 |
| Bond angles (°) | 0.655 |
| Model validation [a] | |
| MolProbity score | 1.59 |
| Clash score | 4.95 |
| Poor rotamers (%) | 1.46 |
| CaBLAM outliers (%) | 2.0 |
| Ramachandran plot quality (%) | |
| Favored | 96.77 |
| Disallowed | 0 |

[a] Assessed using MolProbity (http://molprobity.biochem.duke.edu/).
